# pytom-match-pick: a tophat-transform constraint for automated classification in template matching

**DOI:** 10.1101/2024.09.17.613497

**Authors:** Marten L. Chaillet, Sander Roet, Remco C. Veltkamp, Friedrich Förster

## Abstract

Template matching (TM) in cryo-electron tomography (cryo-ET) enables *in situ* detection and localization of known macromolecules. However, TM faces challenges such as interfering features with a high signal-to-noise ratio and the need for manual curation of results. To address these challenges, we introduce pytom-match-pick, a GPU-accelerated, open-source command line interface for enhanced TM in cryo-ET. Using pytom-match-pick, we first quantify the effects of point spread function (PSF) weighting and show that a tilt-weighted PSF outperforms a binary wedge with a single defocus estimate. We also assess previously introduced background normalization methods for classification performance. This indicates that phase randomization is more effective than spectrum whitening in reducing false positives. Furthermore, a novel application of the tophat transform on score maps, combined with a dual-constraint thresholding strategy, reduces false positives and improves precision. We benchmarked pytom-match-pick on public datasets, demonstrating improved classification and localization of macromolecules like ribosomal subunits and proteasomes that led to fewer artifacts in subtomogram averages. This tool promises to advance visual proteomics by improving the efficiency and accuracy of macromolecule detection in cellular contexts.

## 1 Introduction

Template matching (TM) can be used for the *in situ* detection of macromolecules with known structures in three-dimensional (3D) cryo-electron tomography (cryo-ET) data. TM enables visual proteomics studies, where experimentally determined or predicted structures of macromolecules are mapped to features through the whole volume. Recent advances show that adhering at least to the angular sampling dictated by the Crowther criterion (1), and optimizing spatial sampling and band-pass filters enable confident detection by TM (2). These innovations have become available via GPU implementations that allow sufficient translation and rotation sampling at relevant time scales (1, 3, 4). While TM was shown to outperform recent deep-learning (DL) methods for large assemblies such as fatty acid synthase and the nuclear pore complex (2), annotations from TM nevertheless typically require further manual curation, region of interest (ROI) selection, and/or threshold selection before being used in subtomogram averaging (STA) routines.

To improve detection templates need to be weighted with the cryo-ET point spread function (PSF) (5). This PSF and its corresponding Fourier transform, the 3D contrast transfer function (CTF), have been critical for STA (6, 7), but also improve correlations in TM (4, 8). More elaborately weighted 3D-CTFs model tilt-dependent exposure dampening and CTF parameters, as well as undersampled Fourier space regions due to the tilting scheme. These models additionally require the tomogram to be reconstructed with phase corrections for the CTF and an exposure filter (9, 10). As the metadata for these corrections are not always easily exportable, current methods require adhering to entire pipelines for these steps that keep track of metadata (4, 8). However, the metadata for the PSF in its simplest form only requires a dose and defocus estimate per tilt.

Localization and identification of molecules using TM are often impeded by features in the data with particularly high signal-to-noise ratio (SNR). Such features include strongly scattering contaminations, reconstruction artifacts, or intentionally added gold fiducials for motion registration and targeting. Although the actual shape of these features often varies significantly from the template, their intense scattering results in sharp edges, producing a much higher signal-to-noise ratio (SNR) across all spatial frequencies compared to true positives. True positives display a much lower SNR because biological materials scatter weakly (11). Most TM methods in cryo-EM make use of a locally normalized correlation function which partly accounts for varying contrast throughout a tomogram but does not fully compensate these SNR differences (5, 12).

To our knowledge, two background normalization methods have been suggested to reduce false positives in TM score maps, i.e. the map containing the maximum cross-correlation at each position over the rotational search. Firstly, in 2D TM a whitening filter was suggested, which is calculated as the reciprocal square root of the radially averaged power spectrum of the search image (or volume) (8, 13). This profile is then used as a Fourier filter on both the search image and template, flattening the power spectrum of the image noise and downweighting low-frequency noise that often dominates the signal. Alternatively, STOPGAP introduced simultaneous TM with a phase-randomized template, which correlates the tomogram with a random noise object and thus flattens some background (4). However, both methods reduce correlation with low spatial frequencies and therefore are not suited to fully remove strongly scattering edges that have high SNR over the full range of spatial frequencies. To some extent, the remaining false positives arising from high-contrast artifacts can be filtered out using classification procedures in subsequent STA, but often users are required to manually inspect annotations.

Another issue is that many packages require users to manually specify a correlation threshold to extract positions. Only for relatively strongly correlating, abundant macromolecules, such as ribosomes, the true and false positive rate can be approximately derived from the correlation peak histogram and used for threshold estimation (1). In 2D TM, it has been shown that a threshold can also be estimated by tracking the background variation, with the assumption that the background heavily outnumbers true positives in the position-orientation search space (13). This method is also effective for low-abundance macromolecules.

We introduce pytom-match-pick^i^, an easy-to-integrate open-source command line tool for TM in cryo-ET with GPU acceleration. It supports PSF weighting and different background normalization methods, for which we provide the first quantitative comparison, to our knowledge. To aid automated extraction, we introduce a morphological operation on score maps, the tophat transform, for removing false positives. We show that the transform can be used to estimate a cut-off useful for constraining annotations and automating picking.

## 2 Methods

### 2.1 Tomographic reconstruction

Raw movies of DataverseNL-10.34894/OLYEFI were first motion-corrected using MotionCor2 (1.5.0) (14) without assuming any local motion (due to the low SNR of movies in the tilt series). The frames were already gain-corrected. Corrected frames were prepared for AreTomo (1.3.3) (15) by assembling them into a stack (.st), subsequently, a tilt angle file (.rawtlt) was prepared by extracting the angles from the MDOC file. We also created a text file (.txt) with the total dose accumulation per tilt in e^-^/A^2^ using the same ordering as the tilt angle file. AreTomo was then used to create an aligned stack corrected for local motion assuming 5×5 patches. The alignment was optimized for the tilt axis (starting value -88.7°), sample tilt (but not applied to final reconstruction), and an alignment z-height of 1,000 voxels (corresponding to ∼170 nm). Dose weighting was also applied in AreTomo by adding the accumulated dose to each row of the tilt angle file, and supplying the software with the pixel size and acceleration voltage. IMOD (4.10.29) was used to estimate the defocus for each tilt-series with ctfplotter (amplitude contrast 0.08 and spherical aberration 2.7mm, and 200 keV) (9). It was also used to remove gold beads in the patch-aligned stacks via imodfindbeads and ccderaser. The patch-aligned and gold marker removed tilt-series were then downsampled via Fourier space cropping with IMOD’s newstack by binning 4 times, resulting in a pixel size of 6.9 Å.

Two reconstructions were generated: (i) using CTF-correction with IMOD’s phaseflip followed by AreTomo reconstruction and (ii) using novaCTF^ii^ (10). For (i), strip-based phase flipping was executed with the default minimal defocus tolerance, which dictates strip size, and the tomogram was subsequently reconstructed with weighted-back projection in AreTomo. For (ii), the stacks were CTF-corrected and 3D reconstructed with weighted-back projection in novaCTF, with the estimated defocus parameters and a defocus step of 15 nm. The final tomograms had zyx-dimensions of 500×960×928 at a pixel size of 6.9 Å. Finally, 30 voxels along the x and y edges of the tomogram were tapered using IMOD’s taperoutvol. Tomograms at 8 times binning were also generated from these by Fourier cropping with EMAN2’s e2proc3d.py (2.91) (16).

Reconstruction of *in situ* tilt series from *Chlamydomonas rheinhardtii* cells (EMPIAR-11830) used a similar protocol, except motion-corrected stacks were already available in the repository and could be used directly. Alignment of the tilt series was done without local patches and the gold bead removal step was also skipped. The tilt-axis was optimized from an initial value of -270° in AreTomo. These tomograms were exclusively reconstructed with novaCTF to dimensions 1024×1024×300 voxels and a pixel size of 7.8 Å (4 times binning), and EMAN2’s e2proc3d.py was used to Fourier crop them to 512×512×150 voxels and a pixel size of 15.7 Å (8 times binning).

### 2.2 Calculation of correlation maps

As in our previous work (1), the *LCC* is defined as a function of position x in the tomogram V and rotation u (defined by three Euler angles) of the template T:

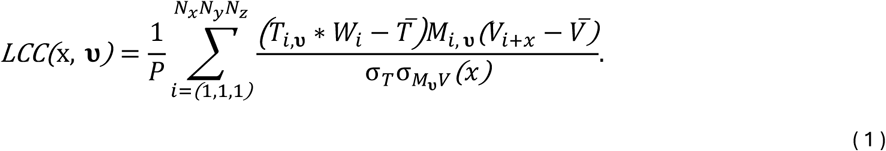

T^-^ and V^-^ are the mean of the template and search volume, T_u_ and M_u_ are the template and mask rotated to υ, σ_T_ is the standard deviation of the template, σ_MuV_ is the local standard deviation of V under M, and P is the sum of the values in the mask. For efficient implementation, we used the Fourier space definition given in (12). The maximum value of the *LCC* at each position x in the search volume is given by:

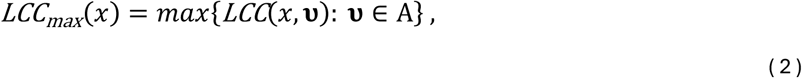

where A is the set of orientations that is searched. For generating angular searches, pytom-match-pick uses the HEALPix package (2024.1) (17) to enable on-the-fly generation of angular searches. This method is commonly used in cryo-ET (18). Additionally, support was added to automatically calculate the angular sampling from the Crowther criterion based on the particle diameter and the provided voxel size or low-pass resolution. By default the TM program assumes a spherical mask, this speeds up computation as M_u_ is identical for each rotation υ, meaning the computationally expensive σ_MuV_ also remains identical. For non-spherical masks a flag can be set to recalculate σ_MuV_ for each rotation of the mask.

The tilt-weighted PSF W is calculated in Fourier space by rotating 2D dose-weighted CTFs to each tilt angle in 3D Fourier space. Tilts are weighted by dose, the B-factor changes by -4 Å^2^ per 1 e^-^/ Å^2^, and the cosine of the tilt angle (7, 8). To account for regions in Fourier space where multiple tilts are overlapping, the CTF for each tilt is also weighted by a ramp filter along the y-axis (tilt-axis) that increases linearly from zero to one at the overlap frequency with the closest neighboring tilt.

### 2.3 Cut-off estimation

The following formula is used to estimate the extraction cut-off τ (13),

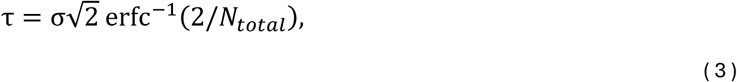

where σ is the standard deviation of the *LCC* over all rotations and voxels, N_total_ is the size of the search space, and erfc^-1^() is the inverse complementary error function. Thus, the cut-off is fully dependent on the search space and the standard deviation. By default, this expression takes the false alarm rate as 1/N_total_, a single false positive over the whole search space. We add a false positive ratio (*FP_ratio_)* parameter that can tune the extraction sensitivity,

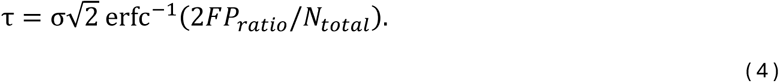

Increasing the FP_ratio_ lowers the cut-off leading to more extracted particles, decreasing it makes the extraction more restrictive.

### 2.4 Tophat transform

Tophat transforms on TM score maps were calculated with the SciPy package (1.13) (19). The ’`white_tophat`’ function from ’`scipy.ndimage`’ was used with a footprint created vià`generate_binary_structure`’. The binary structures were 3D and had a connectivity of 1 unless stated otherwise. The tophat-transformed volume was then sampled on a 1D histogram. A Gaussian distribution was fitted to the log-transform of the histogram and focused only on the first n-bins where the second derivative was larger than zero. Equation (4) was used to calculate a cut-off from this distribution. Particles were extracted if they satisfied both cut-offs.

### 2.5 Background normalization

#### Whitening filter

Normalization with a whitening filter was done by first calculating the radial average of the tomogram’s power spectral density (PSD), where the power spectrum is the absolute square of the Fourier transform. The reciprocal square root of this radial profile was then calculated, *W* = 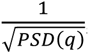 and normalized to a maximum of one to calculate the 1D filter (13). The filter is then interpolated to a 3D spatial frequency grid for the template and tomogram to obtain the 3D Fourier space filter. The tomogram was filtered as a preprocessing step, while the template’s whitening filter was multiplied with the 3D-CTF and thus applied during TM.

#### Phase randomization

For normalization in TM with a phase-randomized object, the random object was first calculated by randomly permuting the phases of the Fourier transform of the template up to the Nyquist frequency. During TM the original and randomized templates are both rotated, convolved with the PSF, and cross-correlated with the tomogram (4). Their *LCC_max_* volumes are separately tracked. Upon completion, the score map of the phase-randomized template is subtracted from the regular score map. The mean value of the phase-randomized *LCC_max_* values is also added to maintain proper background statistics. Particle extraction steps can then be taken as described above.

### 2.6 Manually curated 80S ribosome annotations

To generate ground truth annotations of the 80S ribosome in DataverseNL-10.34894/OLYEFI, TM was first run with a EM map of the human 80S ribosome (EMDB-2938), downsampled to the tomogram’s pixel size, at a 3° angular increment while providing all the necessary data for a tilt-weighted PSF. Particles were first extracted from the TM results using ’`pytom_extract_candidates.py`’ with an exclusion radius of 8 voxels around each peak. The tomograms and the STAR files with annotations were then opened with Blik to inspect and update annotations (20). Only minor changes were needed to remove some false annotations and add missed ribosomes, as judged by eye. The manual annotations are also available in DataverseNL-10.34894/OLYEFI.

### 2.7 Particle classification

#### SHREC’21

The SHREC’21 dataset was used to assess different spatial extents for the kernel used in the tophat transform (21). Specifically, ’`shrec21_full_dataset_no_mirroring.zip`’ was used from the data repository^iii^. For TM, the available atomic models were downloaded and converted to 3D densities using ChimeraX’s molmap (22). Spherical masks were generated for each template that covered the full structure. TM was executed with angular sampling respective to each particle’s diameter. The diameter was chosen along the longest axis for non-globular macromolecules, resulting in the highest possible angle increment for the particle. A binary wedge PSF was used with a defocus estimate of 3.5 μm that was set based on the average defocus range provided in the paper (21). For each particle, we extracted a maximum of 500 candidates with an exclusion radius relative to the particle diameter along its shortest axis, using the default cut-off estimation. For the tophat constraint, the extraction was repeated from the same TM job using different spatial extent for the kernel. All the annotations were then evaluated against the ground truth annotations provided in the SHREC’21 dataset from highest to lowest *LCC_max_* in the list. The FDR, FP/(TP+FP), and recall, TP/(TP+FN), were calculated as a function of *LCC_max_*. The point in the curve where (recall * (1-FDR)) was highest, was taken as the rectangle under the curve (RUC).

#### DataverseNL-10.34894/OLYEFI

(This is an excluded set of EMPIAR-11830.) Localization of 60S ribosomes against manually curated 80S ribosome annotations was done to assess multiple TM enhancements. The 60S ribosome template was generated from an atomic structure of the human 80S ribosome (PDB 6qzp) by using large subunit labeled chains. For testing the effects of the binary wedge PSF the tilt angle file was provided with a single defocus estimate (3 μm), while for the tilt-weighted PSF the ’`--per-tilt-weighting`’ flag was set together with a text file with tilt-dependent defocus estimates (.defocus) and a text file with the dose accumulation (.txt). The angular sampling increment for TM was set to 3°.

For both background normalization methods, the tilt-weighted PSF was used together with a 3° angular sampling. The whitening filter can be set with the ’`--spectral-whitening`’ flag while phase randomization is set with the ’`--random-phase-correction`’ flag in ’`pytom_match_template.py`’. To compare the effects of background normalization, annotations were extracted from a baseline job and jobs with either of the normalization methods activated. All jobs were forced to extract 500 particles. Similarly to the analysis in the SHREC dataset, the annotations were compared against the curated 80S annotations to analyze the ROC and determine the RUC value.

To compare the effects of the tophat constraint, annotations were extracted from the phase randomization job with and without the ’`--tophat-filter`’ flag while keeping the kernel connectivity at 1, i.e. the kernel with the least spatial extent. For both the baseline and tophat constraint the *FP_ratio_* parameter was kept at its default of 1, and particles were extracted up to the determined thresholds. The recall and precision (TP / (TP+FP)) were calculated only on the full list— not as a function of the *LCC_max_*. The f1-score is calculated as the harmonic mean of precision and recall: 2 * (precision * recall) / (precision + recall).

#### EMPIAR-11830

26S proteasome localization was performed in 8x binned tomograms reconstructed with novaCTF. For the template, an SPA reconstruction of the yeast proteasome (EMD-6575) was recentered on one of the regulatory caps of the 26S proteasome and then downsampled to 15.7 Å. A mask was created to cover the regulatory particle and 50% of the 20S core. TM was done with the tilt-weighted PSF and an angular sampling of 7°. Particles were annotated with and without the tophat constraint to compare the effects of the filter.

### 2.8 STA

Averaging of 60S ribosomes was done as described in Chaillet*, et al.* (1). Specifically, subtomogram were reconstructed at a voxel size of 3.45 Å and a box size of 120 voxels, wide enough for the reconstruction to cover the small ribosomal subunit and the ER-membrane. Local resolution estimation after refinements was done using RELION’s built-in implementation.

Dynamo was used for averaging 26S proteasomes (1.1.532) (23). Dynamo2m was used to convert STAR files with particle annotations to dynamo tables. The dynamo tables were used to crop subtomogram from the 4x binned reconstruction of novaCTF, using the dynamo software. These were then aligned and averaged for four iterations using the dcp graphical user interface. Aligned particles were displayed on a tomographic slice using ArtiaX.

### 2.9 Python software packages

Although not mentioned explicitly, these scientific Python packages have also been essential for this work: mrcfile (24), starfile (25), NumPy (26), voltools^iv^, Matplotlib (27), seaborn (28).

## 3 Results

### 3.1 pytom-match-pick is a GPU-accelerated TM module that is easy to integrate

pytom-match-pick is a command line interface (CLI) Python package for GPU-accelerated TM in tomograms (1). The package aims at ease of use while maintaining a simple codebase. Under the hood, pytom-match-pick calculates the maximum of the local correlation coefficient (*LCC*) of the tomogram and the template over the rotational search space, which we refer to as the *LCC_max_* (Equation (2) in Methods). For each rotation, the template is interpolated and convoluted with the PSF before computing the FLCF. Data for the PSF model is extendable based on available metadata: for example, if dose weighting is not to be used, a binary missing wedge in Fourier space can be specified by providing tilt angle limits while providing tilt angles in the IMOD ‘.rawtlt’ or ‘.tlt’ format allows per-tilt weighting (Figure S1 A; left) (9). Additional defocus files (IMOD’s ‘.defocus’) and accumulated dose (e^-^/A^2^ per line in ‘.txt’) can be provided for CTF and dose-weighting (Figure S1 A; right). The output is written as a STAR format particle list. This enables visualizing the annotations in the napari-based (29) viewer Blik (20), while results can be used for STA in RELION (18) or Warp (8).

### 3.2 pytom-match-pick has improved performance and template weighting

pytom-match-pick has improved speed compared to our previous implementation (1). Real-spaced input FFT routines in Cupy (30) that make use of built-in caching for recurrent calls enable a roughly 2-fold speed-up compared to PyTOM (Table S1).

Furthermore, we implemented the tilt-weighted PSF and compared its performance to a binary-wedge PSF with a single defocus estimate using the example of 80S ribosomes associated with vesicles derived from the endoplasmic reticulum (31). This is also the pytom-match-pick tutorial dataset, which we here refer to as DataverseNL-10.34894/OLYEFI^v^. For the comparison we reconstructed the tilt series with two different approaches to CTF correction: IMOD’s ctfphaseflip (9), which accounts for the CTF gradient in tilted projections, and novaCTF (10), which additionally accounts for the defocus gradient within the reconstructed volume. A visualization of the two PSFs illustrates that the tilt-weighted PSF models all the undersampled regions in Fourier space and a tilt and dose-dependent drop-off (Figure S1). Additionally, the amount of ‘fanning’ of the PSF is dependent on the number of sampling points in Fourier space, which is, in turn, a function of the box size of the template (Figure S1 A). Both, the tilt-dependent weighting of the template and reconstruction with novaCTF increase the correlation coefficients of particles, making them more easily distinguishable from the background, indicated by the increased *LCC w*hen normalized against standard deviation (σ) (Figure S1 B). Reconstructing the tomogram with novaCTF gives the most notable improvement compared to the baseline (binary PSF and ctfphaseflip). In addition, the benefit of a tilt-weighted PSF is more pronounced in conjunction with novaCTF. These calculations were performed in tomograms with a maximal resolution (Nyquist) of (28 Å)^-1^ (14 Å voxel size), and the effects might become more pronounced in reconstructions with increased sampling. Overall, the combination of detailed PSF modeling and accommodating defocus gradients in the reconstruction improves the correlations in TM.

### 3.3 Phase randomization improves 60S ribosome classification

We then compared previously suggested background normalization methods for the classification of the ribosomal large 60S subunit in DataverseNL-10.34894/OLYEFI. Due to its lower molecular weight, the false positive rate is expected to be substantially larger for the 60S subunit compared to the search for the full 80S ribosome. Both spectrum whitening (13) and phase randomization (4) were implemented. Phase randomization is computationally more expensive because the local correlation also needs to be evaluated for the phase-randomized version of the template, while spectrum whitening only modifies the constant weights leading to a marginal increase in preprocessing time. We first ran TM with the 80S ribosome with a 3° angular increment, and manually curated annotations to obtain a ‘ground truth’ list (available in the DataverseNL repository). Classification statistics of TM with the 60S ribosome could then be compared with these annotations (Figure 1A).

**Figure 1.**
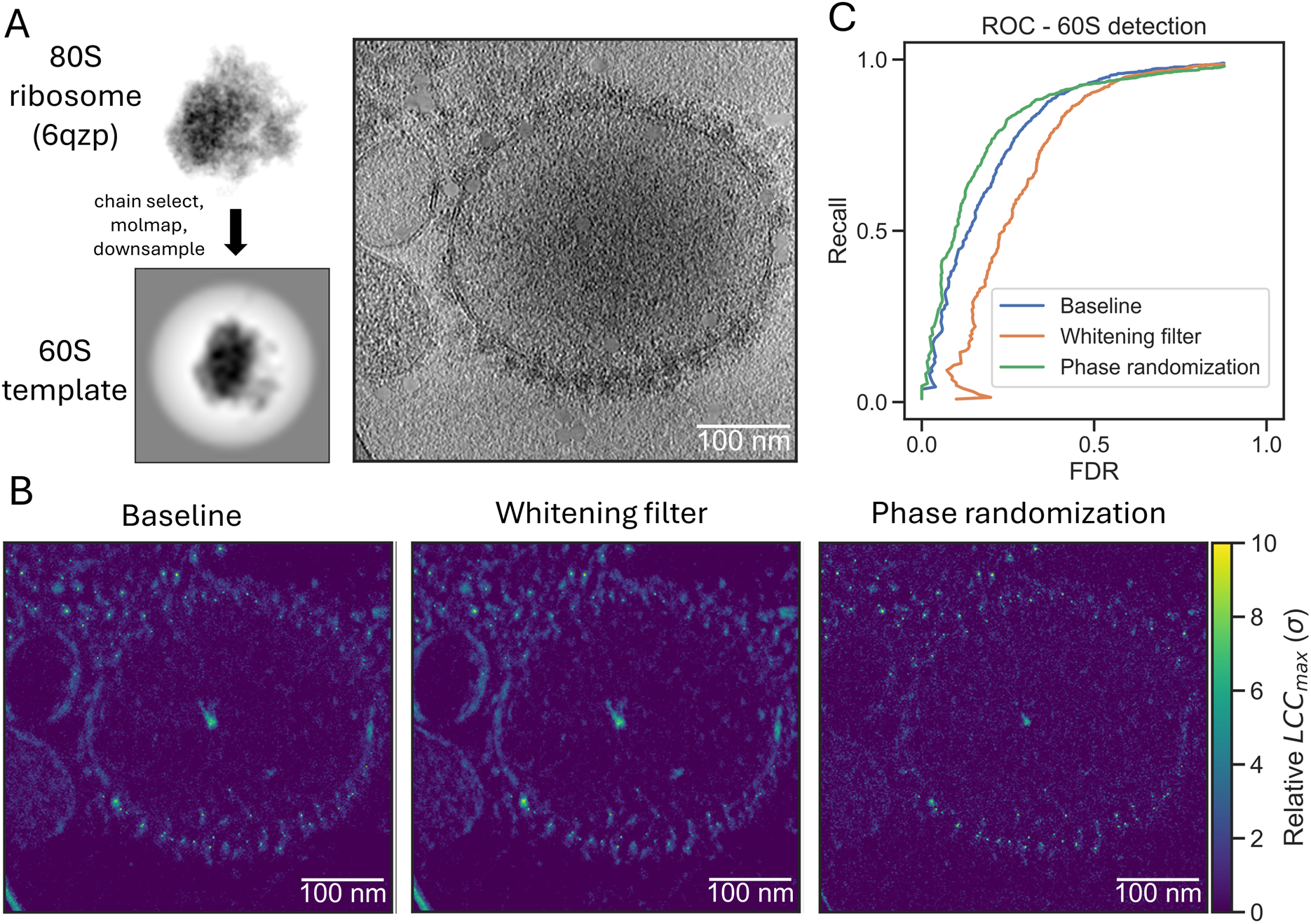
Background normalization improves TM-based identification for 60S ribosomes. (**A**) An example of the required input for TM in pytom-match-pick. An atomic model of the 80S ribosome can be used as input from a database. Specific protein/RNA chains can be selected from the model. It needs to be sampled to the spacing of the tomogram and a mask needs to be generated. A tomogram of ER-derived vesicles is used to detect the structure in 3D. (**B**) Illustration of TM score maps (*LCC_max_*) with the 60S large ribosomal subunit for the baseline method (left), with a whitening filter applied (center) and with flattening via phase randomization (right). The images are maximum projections of the 3D score maps along the z-axis. (**C**) A plot showing the ROC of 60S ribosome detection for the baseline (blue) and two background normalization methods (whitening filter – orange; phase randomization – green). The FDR and recall were calculated over 10 tomograms by comparing them against curated annotations.

In the TM score maps the whitening filter enhances sharp edges and membranes, while phase randomization reduces these features compared to the baseline (Figure 1B). The receiver-operator curves (ROC), which plot the recall against the false discovery rate (FDR), show that phase randomization slightly improves classifier performance, while the whitening filter reduces performance compared to baseline TM (Figure 1C). While all methods level at approximately the same recall, the FDR is increased for the whitening filter leading to worse identification. In contrast, phase randomization reduces the number of false positives.

### 3.4 A tophat transform can separate peaks and artifacts

We next examined whether morphological operations could further improve annotation in TM. First, we noted that high-contrast false positives often produce extended patches of high values in score maps, while true positives tend to show steep local maxima (i.e. peaks) in the correlation map (Figure 2A) (2, 8, 13). We reasoned that a transformation that can separate extended patches from peaks would be appropriate to improve specificity. Hence, we tested a tophat transform, which is effective at detecting speckles in astronomy images. While the purpose in astronomy is to remove fine features from images, we aimed at the opposite, i.e., filtering out larger features using the tophat transform. Figure 2B illustrates the effectiveness of this transformation in separating a sharp peak from the background based on a defined small kernel. In TM score maps the tophat transform removes large patches (Figure 2A).

**Figure 2.**
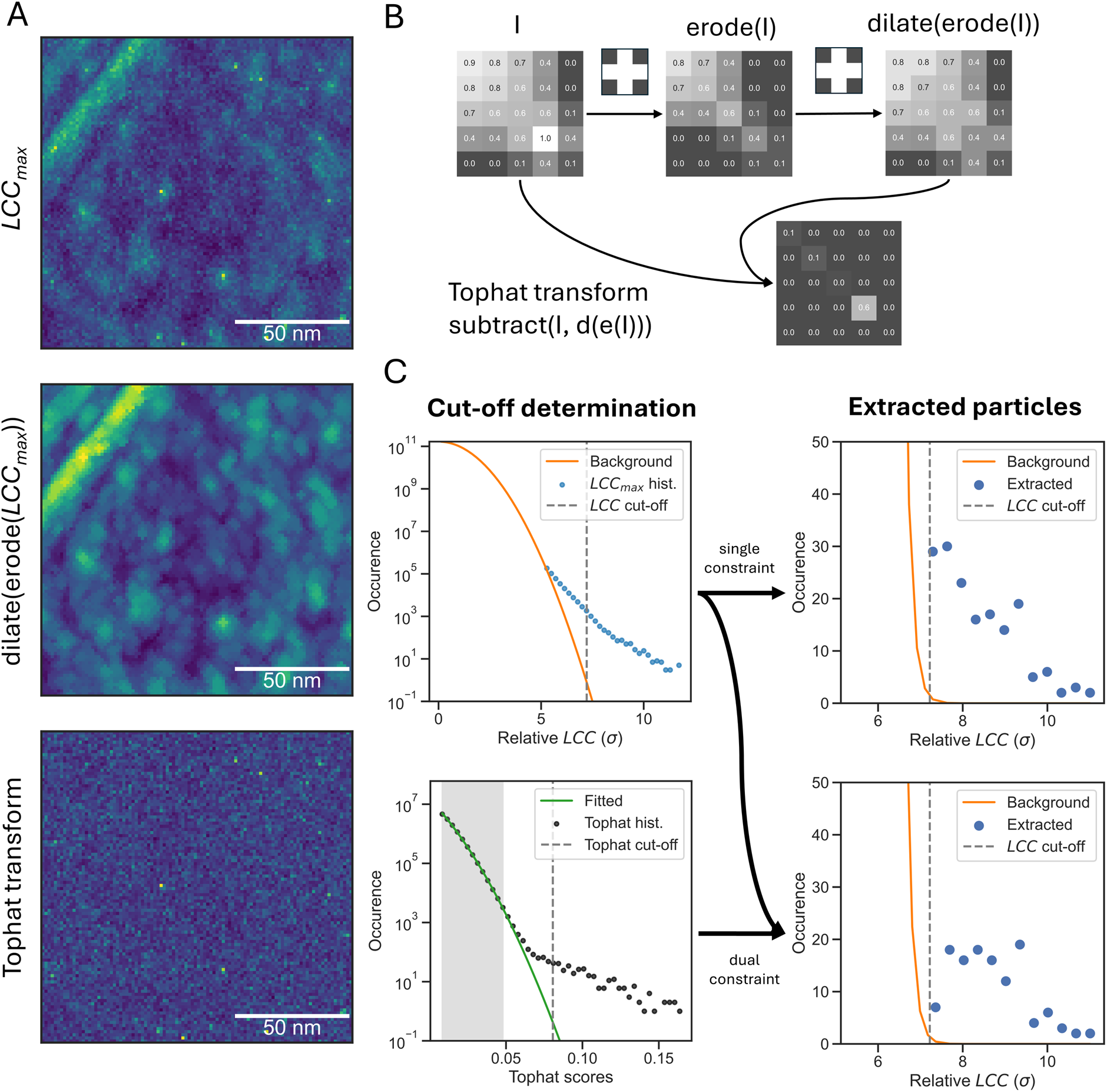
A tophat transform can reduce background and automate TM annotation. (**A**) Maximum intensity projections along the z-axis of a score map of 60S LSUs matched with microsome tomograms (top), the morphological opening of the score map (erosion followed by dilation, center), and a tophat transform of the score map (bottom). (**B**) A 2D illustration of a tophat transform on a small image with intensity values annotated in each pixel. The tophat transform involves an erosion followed by a dilation within the specified kernel. The tophat transform is then obtained by subtracting this morphological opening from the original image. (**C**) Automated cut-off estimation in pytom-match-pick. The top left plot shows the default cut-off estimation (see Methods), which determines a threshold (grey dashed line) based on the Gaussian distribution of *LCC* from TM (orange line). The blue dots indicate *LCC_max_* values above the background. The top right shows the distribution of extracted particles after applying the cut-off according to the error function (Equation (3)) and filtering peaks. The bottom left shows the cut-off estimation (dashed line) based on the fitted Gaussian (green) on a histogram of the tophat transform (black dots). Both cut-offs can be simultaneously applied to obtain the final extracted particles that are above both cut-offs (bottom-right).

### 3.5 Dual constraint cut-off estimation automates annotation

We examined if we could automatically extract candidates using the thresholding approach introduced by Rickgauer, Grigorieff and Denk (13) for 2D template matching and combine it with the tophat transform. To estimate a threshold, Gaussian background noise is assumed to greatly outnumber the true positives. For background noise estimation, the square sum of the *LCC* values is accumulated during the rotation search. The variance of the *LCC* values can then be obtained by dividing this sum by the full search space (*N_total_ = N_voxels_ × N_rotations_*). This procedure provides the variance of the *LCC* values over the whole search with an expected mean of zero (Figure 2C; top-left). A score threshold can be determined from this distribution, as the error function over this distribution returns an expected number of false positives. pytom-match-pick applies this threshold by default on extraction, but the expected false positives can be reduced or increased for less or more sensitivity. We further refer to this peak selection as the ‘baseline’ method.

We then plot the distribution of scores after extracting particles based on the determined cut-off. As extraction takes into account the particle diameter to prevent double annotation of the same particle in a tomogram, the final distribution is altered (Figure 2C; top right). Therefore, the occurrence of scores is heavily reduced after extracting particles (Figure 2C; top row).

We applied a similar method for finding a threshold on the tophat transform of the *LCC_max_* values. Although it is not feasible to track the variance during the rotational search, a histogram of the transform shows a clear Gaussian background distribution (Figure 2C; bottom left). Based on the log-transform of this histogram, a Gaussian is fitted to the first *N* points where the second derivative is larger than zero. A threshold is set similarly to the *LCC_max_* values, i.e., based on the expected false alarm rate. Both the *LCC_max_* and tophat transform threshold are used as a dual constraint for extracting particles, which we further refer to as ‘tophat constraint’. The resulting *LCC_max_* values of the extracted particles—a union of both thresholds—have reduced occurrence as the tophat transform constrains the annotations further (Figure 2C; bottom right).

### 3.6 A spatially restricted kernel for the tophat transform improves classification in SHREC’21

We benchmarked the annotation for the baseline and tophat constraint methods on the SHREC’21 dataset. We furthermore evaluated the influence of the spatial extent of the kernel in the tophat transform using this simulated dataset. Three kernels with increasing spatial extent were examined (Figure 3). For the four candidates (baseline and 3x tophat), we calculated ROC curves against the ground truth annotations in the SHREC’21 evaluation tomogram (Figure S2). We note that in some cases the ROC curve has data points with an FDR close one and a low recall, which occurs if the highest-ranking annotation is a false positive such as a gold bead (Figure S2). Among the different kernels tested the one with the smallest spatial extent (connectivity of 1) performs best (Figure 3B). With increasing spatial extent of the kernel the ROC approaches the baseline performance. Thus, this limited comparison suggests that the tophat transform is most effective with a small spatial extent when it filters for sharp peaks in the score map most aggressively.

**Figure 3.**
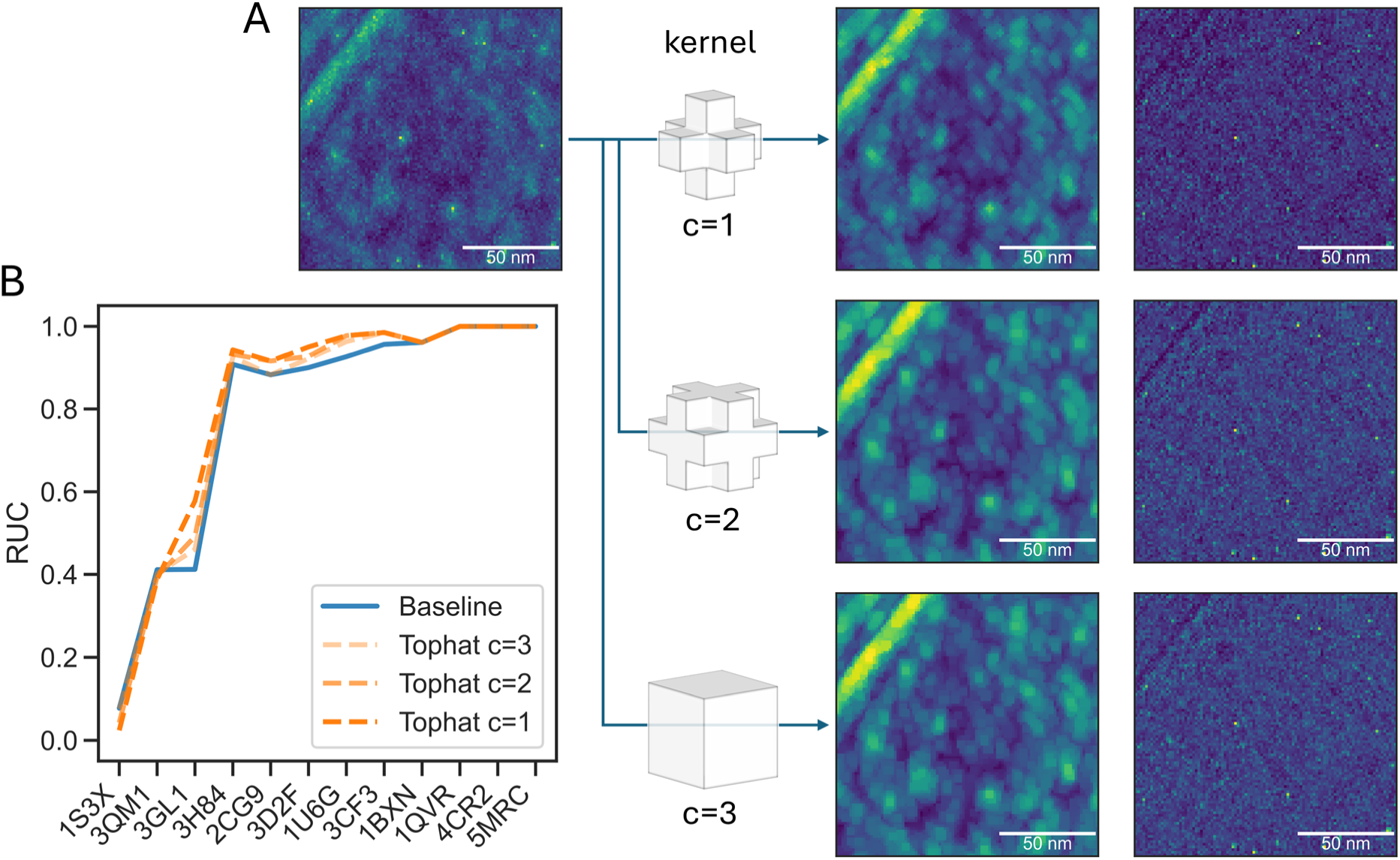
A tight kernel for the tophat transform provides the best classification in SHREC’21. (**A**) Visual inspection of the tophat transform using kernels with different spatial extents (or connectivity, denoted by ‘c’ in the figure). The kernels are shown together with their greyscale opening and the final tophat transform. (**B**) Classification results with increasing connectivity (dark to light orange dashed lines) in SHREC’21 compared against the baseline (blue line). The RUC is shown per ground truth particle in the dataset where the particles are ordered by increasing molecular weight from left to right.

### 3.7 Matching 60S ribosomal subunits with the tophat constraint improves precision

We further compared the performance of the tophat transform to the baseline method by employing baited reconstruction (32, 33). Here, the search results of the 60S subunit serve as bait for the reconstruction of the complete 80S ribosome from the corresponding subtomograms. Using only this limited bait (Figure 1A), we can assess possible reference bias based on expected structural features that are absent from the template and should appear upon STA, such as the 40S ribosomal subunit and the ER membrane, similar to the originally reported structure (31). Visual inspection of the annotations in a single tomogram shows that the union of the tophat and baseline annotations is mainly clustered around an ER-derived vesicle, while the unique annotations in the baseline method are spread around the ice (Figure 4A). The comparison of the 60S detections with the manually inspected 80S annotations in 10 representative tomograms indicates that the precision improves substantially upon tophat filtering at the expense of a small decrease in recall (Figure 4B). An improvement in the overall performance is reflected in the f1-score, which is the geometric average of recall and precision (Figure 4C).

**Figure 4.**
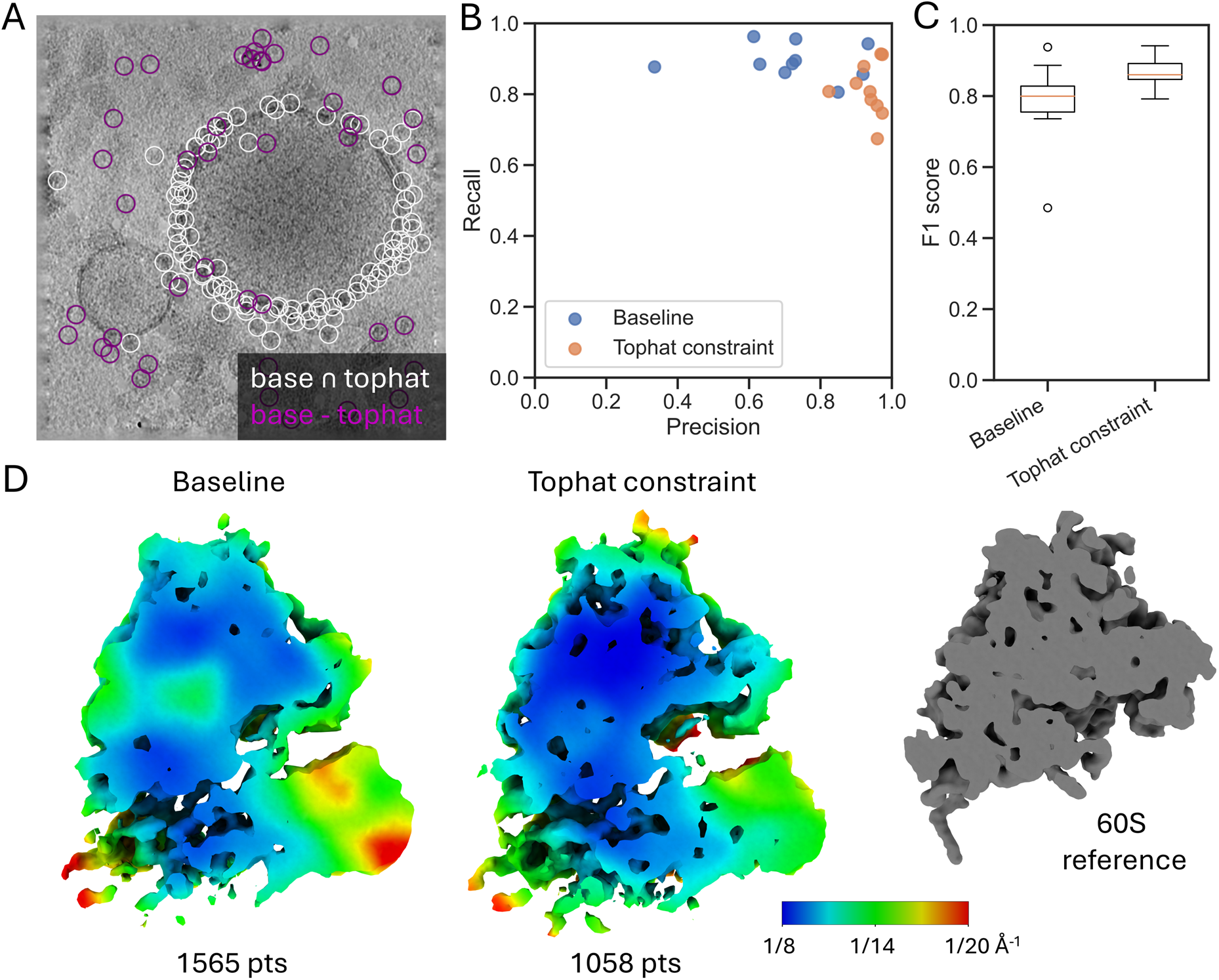
The tophat constraint improves 60S ribosome annotation in experimental data. (**A**) Union (white) and exclusion (purple) of the baseline and tophat constraint annotations in a single tomogram from the DataverseNL-10.34894/OLYEFI. The image is a projection of the tomogram along the z-axis and the circles are also projected from the 3D annotations onto the image. (**B**) Precision and recall of the test dataset for the baseline (blue dots) and tophat constraint (orange dots) annotations, each dot represents the precision/recall on a single tomogram. (**C**) A box plot of the f1-scores per tomogram (N=10) for the baseline and tophat constraint, where dots indicate points falling outside 1.5 times the interquartile range. (**D**) Local resolution of reconstructions from STA (after alignment). Both methods show structures not present in the original 60S reference (grey structure on the right). The left image shows the local resolution of the baseline set (1565 particles); the right image show the local resolution of the tophat constraint set (1058 particles). The colorbar ranges from (8 Å)^-1^ (blue) to (20 Å)^-1^ (red). The number of particles in each set is noted below the image.

To further estimate the quality of both sets, we used STA to resolve the structure and estimated the local resolution. Although both sets show an average containing both the 60S and 40S subunits (confirming our baited reconstruction check), the average derived from the baseline set reaches a maximal local resolution of (8.5 Å)^-1^, while the tophat-constrained set reaches (7.9 Å)^-1^ with only 2/3 the number of particles (Figure 4D). Additionally, the tophat-constrained set reaches higher resolution throughout the stable 60S subunit and part of the 40S subunits. This confirms the reduction of false positives in the tophat-contained set. Overall, the tophat constraint improves the quality of the multi-particle reconstruction.

### 3.8 Proteasome bait in Chlamydomonas retrieves 26S proteasome with the tophat constraint

To further characterize the performance of the tophat constraint we used TM to localize a part of the 26S proteasome in a recently released dataset of *Chlamydomonas rheinhardtii* cells (34). Similarly to the ribosome bait, a section of a 26S proteasome, reconstructed with STA, (EMDB-3932) was used as the template by masking (Figure 5A). The bait contained 50% of the 20S core particle and one regulatory particle. True positives should all display the full core particle and many also a second regulatory particle associated with it. Three tomograms recorded in the vicinity of the nuclear envelope and ER were used, where proteasomes are likely to localize (35). After TM, the proteasome annotations of the baseline and tophat constraint resulted in 100 and 32 particles, respectively. STA of the baseline set gave rise to a structure resembling a 26S proteasome but with clear artifacts, while an STA of the tophat-constrained set produced a structure resembling the full 26S proteasome where the 20S core particle and a second regulatory particle is resolved (Figure 5B).

**Figure 5.**
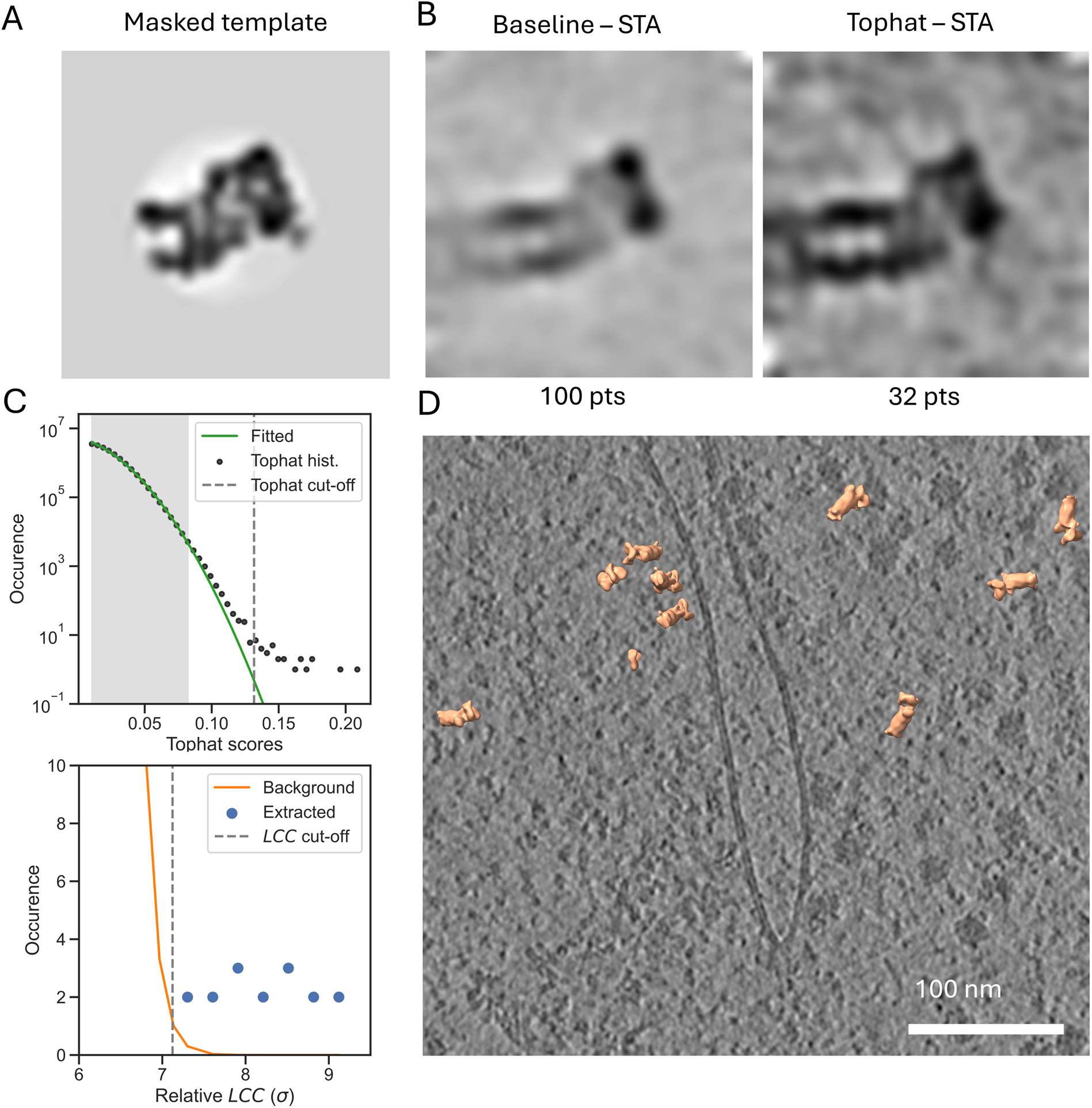
The tophat constraint improves 26S proteasome classification for in situ lamellae. (**A**) The center slice of the template after multiplication with the generated spherical mask. It illustrates what part of the original structure is removed. (**B**) Center slices of the reconstruction via STA (after alignment) of the baseline set (100 particles) and tophat constraint set (32 particles). Both show recovered structures not present in the initial template. The number of particles in each set is displayed below. (**C**) The top plot shows the Gaussian that was fitted (green line) to the histogram of the tophat transform values (black dots) and the corresponding cut-off (dashed grey line). For fitting the Gaussian only the gray shaded area was used as those points satisfied a positive second derivative. The y-axis shows the log of the occurrence of each bin. The lower plot shows a histogram of the *LCC* (normalized by σ) occurrence of the annotated particles (blue dots) after the tophat constraint. (**D**) A cropped slice of a tomogram focused on a nuclear pore. Particles annotated by the tophat constraint are visualized on top of the slice.

Based on the STA and the statistics for the cut-off estimation (Figure 5C), the tophat constraint is relevant for a macromolecule that is of markedly lower abundance than the ribosome. Inspecting the proteasome annotations in an abundant section of a tomogram around a nuclear pore, some proteasomes localize close to the pore within the nucleus (Figure 5D), consistent with previous work (35). TM and the tophat constraint are applicable for automated localization of a low abundance ∼1 MDa complex in full tomograms.

## 4 Discussion

### 4.1 Closely matching the tomogram and template via PSF modeling

A close match between the target macromolecule in the tomogram and the reference is important for TM. We and others previously showed that angular sampling is a critical parameter determined by the particle diameter and resolution that is needed for accurate matching (1, 2). Furthermore, for a correct match, the CTF needs to be accounted for in the reconstruction of the tomogram and the weighting of the template. We showed that for the reconstruction the CTF correction using novaCTF is more effective than strip-based phase flipping in IMOD for TM-based localization. For the template, a tilt-weighted PSF model is slightly more effective. Such improvements have also been previously demonstrated by others (4, 8). We expect that this becomes more important at smaller pixel sizes (increased spatial sampling) of the reconstruction and for smaller macromolecules. Although these steps improve performance, we emphasize that a binary wedge PSF with a single defocus estimate can already be efficient at classifying large structures (1, 3).

Wan, Khavnekar and Wagner (4) also suggest using cisTEM’s electrostatic potential calculation (36) instead of ChimeraX’s molmap (22). The difference in these methods is the parameterization of the scattering factors: Chimera uses a single Gaussian per atom, cisTEM uses a sum of 5 Gaussians parameterized from quantum chemical calculations as commonly used for atomic modeling into EM maps (37). Additionally, cisTEM can use a model for solvent exclusion that likely enhances the simulation of the electrostatic potential. However, it has not been quantified to what extent these effects influence the match of the reference with the tomogram. It is generally known in the field that using previously determined maps from the EMDB or STA of subsets is most effective for localizing targets. It would be interesting to see how well simulated electrostatic potentials could approximate results with an EM-reconstructed reference.

### 4.2 Phase randomization improves particle classification

We quantified the effects of a spectrum whitening filter and noise flattening via phase randomization to reduce the response to strongly scattering background objects. In our analysis, phase randomization seems a reliable method for TM that reduced the response to false positives, as indicated by the decreased FDR, spectrum whitening on the other hand increased the FDR. Earlier results suggested that the whitening filter in tomograms resulted in steeper local maxima for true positives (8), however, it was not evaluated for classification performance, which we did here. In comparison with 2D TM, where the whitening filtering was reported to improve classification (13), the poorer classification performance in tomograms might be due to the reduced SNR in the high-frequency regime compared to 2D data acquired with the entire allowed electron dose. Additionally, in 2D TM, the whitening filter was shown to be effective due to the strong weighting of the high-resolution signal by working with low defocus values. However, data acquisition at low defocus values might not be as beneficial in cryo-ET due to the increased difficulty of aligning the tilt series to a common coordinate system. Our results suggest that spectrum whitening might reduce the classification performance of TM in tomograms, while normalization of the score map via correlation with a phase-randomized template likely improves classification (4).

The phase randomization method might be further expanded by randomizing the phases only beyond certain frequency shells. This is commonly used for the estimation of mask bias in cryo-EM for Fourier shell correlations (38). However, in TM, it would be similar to setting a high-pass filter which was already extensively evaluated by Cruz-Leon*, et al.* (2), while using the phase randomization method in STOPGAP. From their study, it seems dependent on the type of macromolecule: 80S ribosomes produce the highest average peaks when including all low-spatial frequencies, while microtubules and ribosomal vaults can benefit from removing frequencies up to (200 Å)^-1^. Thus, for certain macromolecules, it might be beneficial to remove some low-spatial frequencies.

### 4.3 Morphological operations can be effective in removing false positives

Morphological operations can be an efficient tool to post-process TM results. We illustrate this with the tophat transform that distinguishes sharp peaks in the score map and constrains annotations to local maxima. Similarly, Balyschew*, et al.* (39) recently proposed post-processing score maps by removing large islands of connected components. However, they did not show to what extent this improves classification by TM. Our results on the SHREC’21 dataset and the ribosomes on ER microsomes show detailed classification statistics that show such methods can be effective.

A downside of these methods is that an additional filtering step by definition reduces particle recall. Such trade-offs can be worthwhile if they improve overall classifier performance. Especially for STA procedures, where specificity is more important than recall, they can save on precious manual labor. If maximal sensitivity is desired, it might not be worthwhile if methods in downstream processing can safely remove false positives.

### 4.4 Using autocorrelation functions

Although the tophat transform is effective for score map filtering, it only exploits the spatial extent of the *LCC_max_*. A more comprehensive approach would be to exploit both the spatial and rotational aspects of the correlation function. The cross-correlation scores for a given template have a unique spatial and rotational structure defined by its autocorrelation function (2). Rather than filtering for steep local maxima, using the autocorrelation function could provide a more precise method for constraining annotations. However, as it is a function of translation and rotation, tracking autocorrelation functions for analysis will require some solutions to work around this multidimensional data.

### 4.5 Conclusions

Here, we introduced pytom-match-pick, an open-source GPU-accelerated tool for TM in cryo-ET. We demonstrate improvements in TM through tilt-dependent PSF modeling and background normalization techniques. Notably, we introduce a novel approach using a tophat transform to reduce false positives and automate annotation. This method shows particular promise for detecting lower abundance macromolecules like the 26S proteasome in cellular contexts. Overall, pytom-match-pick offers an accessible and effective solution for researchers aiming to locate and identify macromolecules in cryo-ET data, potentially advancing *in situ* structural biology.

## Data availability

The data for this study was downloaded from DataverseNL, EMPIAR, EMBD, and PDB with identifiers mentioned in the text.

## Code availability

pytom-match-pick is available on GitHub via: https://github.com/SBC-Utrecht/pytom-match-pick.

## Author contributions

Conceptualization, M.L.C., R.C.V, and F.F.; methodology, M.L.C., S.R., R.C.V, and F.F.; software, M.L.C. and S.R.; validation, M.L.C.; formal analysis, M.L.C.; investigation, M.L.C.; resources, F.F.; data curation, M.L.C.; writing—original draft preparation, M.L.C. and F.F.; writing—review and editing, all authors; visualization, M.L.C.; supervision, R.C.V, F.F.; project administration, F.F.; funding acquisition, F.F. All authors have read and agreed to the published version of the manuscript.

## Declaration of generative AI in the writing process

During the preparation of this work the authors used ChatGPT in order to improve paragraph readability. After using this tool/service, the authors reviewed and edited the content as needed and take full responsibility for the content of the published article.

## Acknowledgments

We thank Hamid Rahmani for suggestions on tomogram masking and Ricardo Righetto and Philippe van der Stappen for the discussion on implementing a tilt-weighted PSF and continuous feedback.

## Footnotes

i https://github.com/SBC-Utrecht/pytom-match-pick

ii Commit 4f134c7 on branch ‘master’; https://github.com/turonova/novaCTF.

iii https://doi.org/10.34894/XRTJMA

iv https://github.com/the-lay/voltools

v https://doi.org/10.34894/OLYEFI, DataverseNL, V1

