## Supplementary Data for "pytom-match-pick: a tophat-transform constraint for automated classification in template matching"

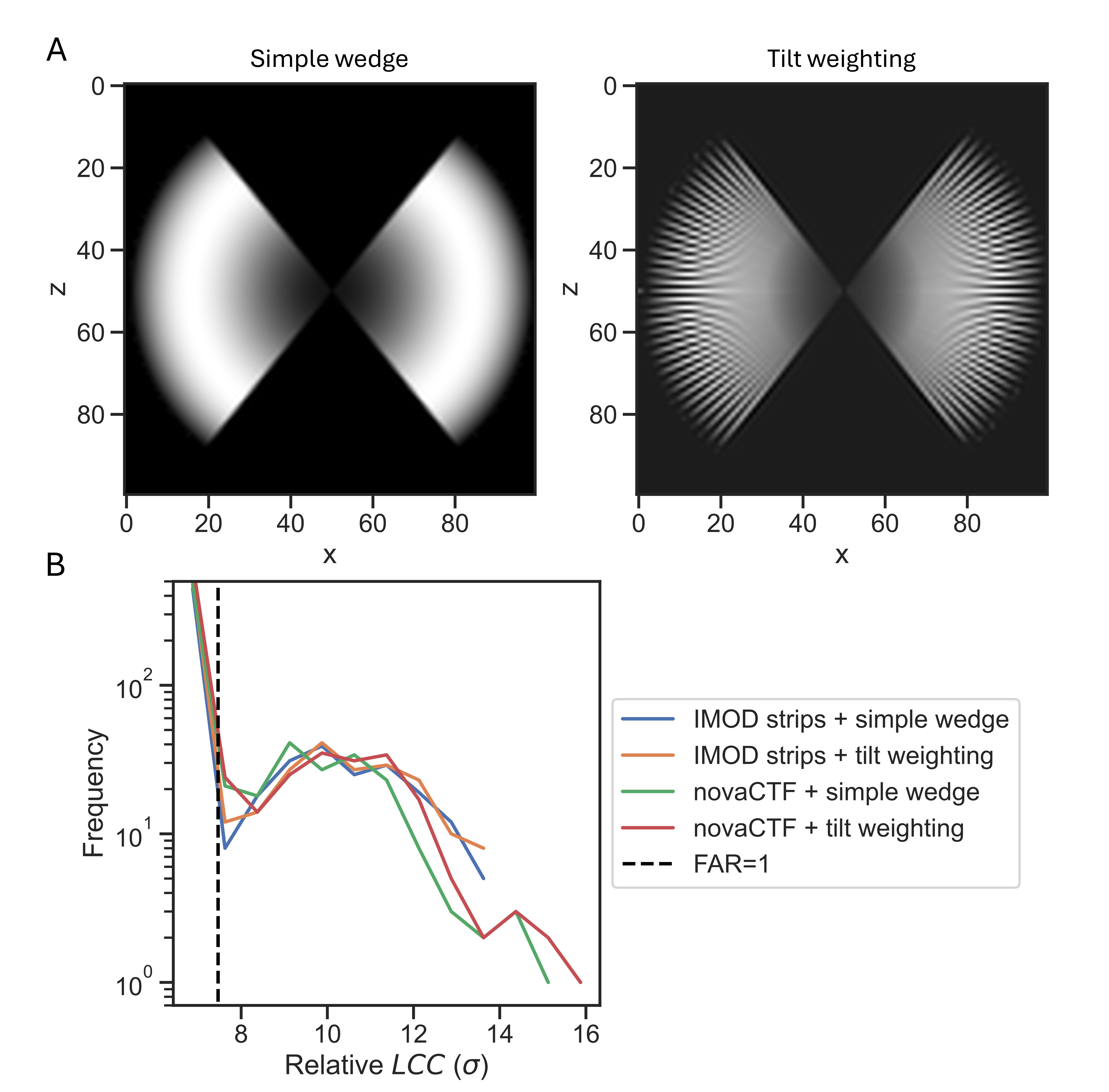

### Figures

Figure S1. **3D-CTF models for the tomogram and template improve the cross-correlation.** (**A**) An illustration of two PSF models in Fourier space: the left image shows a binary wedge model with single defocus for the whole tilt-series (simple wedge), and the right shows a weighting scheme that includes tilt-dependent CTFs and exposure dampening (tilt-weighting). Both models are shown as a slice along the y-axis (the tilt-axis). (**B**) Effects of different reconstruction methods and PSF in TM. Each line shows a histogram of the occurrence of the LCC_max_ of an extracted particle list after division by the standard deviation, σ, as tracked during TM. The blue and orange lines are IMOD reconstructions with the simple and tilt-weighting from (A), respectively. The green and red lines are novaCTF reconstructions with a binary wedge PSF and a tilt-weighted PSF, respectively. The dashed vertical line indicates the cut-off with an expected FAR (= false alarm rate) of 1.

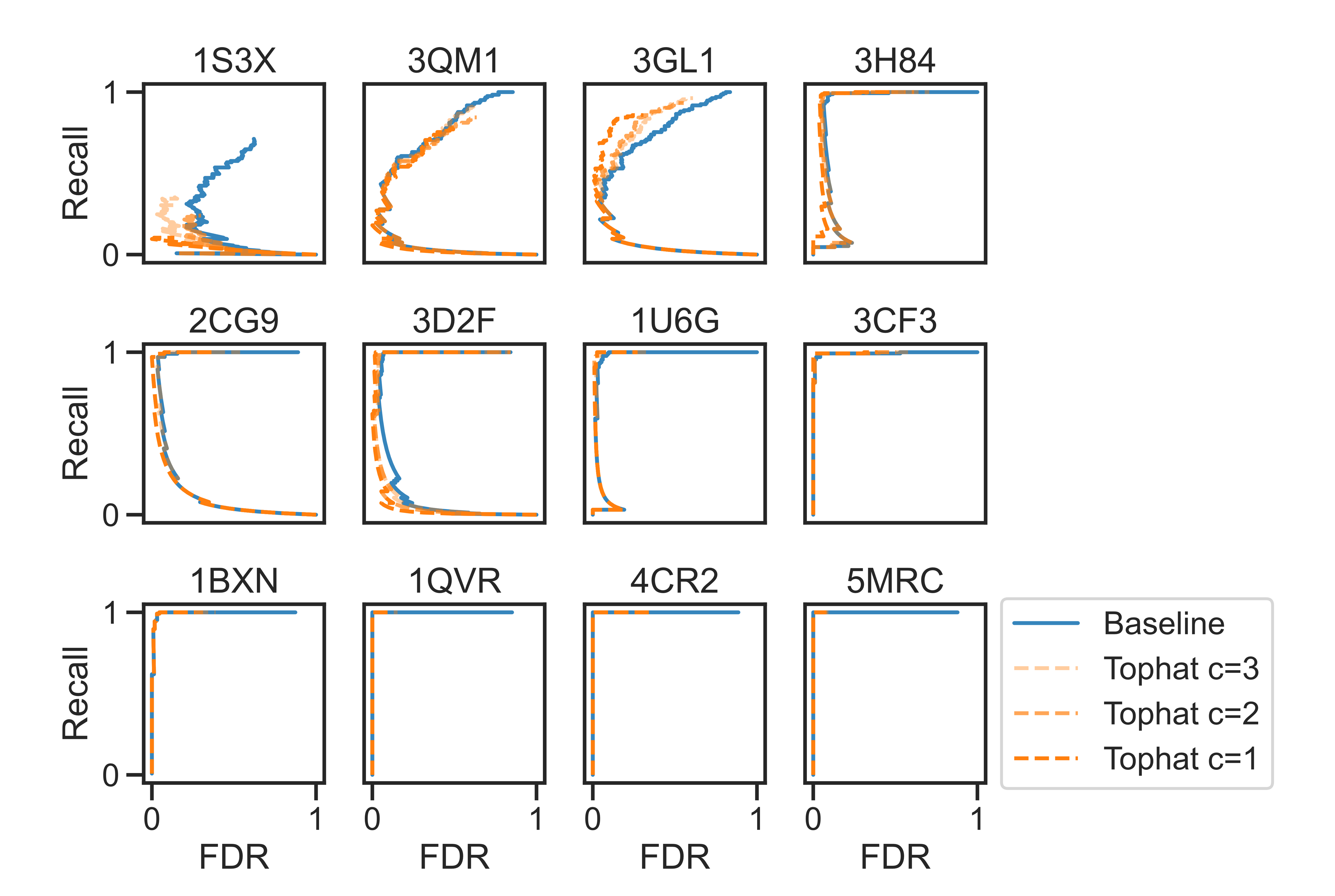
Figure S2. **Per particle ROC for the SHREC’21 evaluation tomogram.** Each plot shows four ROCs for the baseline (blue line), and tophat with increasing spatial extents (connectivity) (dark to light orange dashed lines). The x-axis shows the FDR and the y-axis the recall, the panel titles indicate the PDB ID of each particle.

### Tables

Table S1. **pytom-match-pick has improved performance.** Timing was compared against our reported timings in Chaillet et al. (2023) on identical hardware and settings.

|  | Setup | Timing |
| --- | --- | --- |
| PyTOM (Chaillet et al., 2023) | 7° sampling, 4 GTX 1080 Ti | 661 s. (11.0 min.) |
| pytom-match-pick | 7° sampling, 4 GTX 1080 Ti | 306 s. (5.10 min.) |
